# Brain virtual histology of a lizard species (*Podarcis bocagei*) using X-ray micro-tomography and deep-learning segmentation

**DOI:** 10.1101/2024.07.05.602071

**Authors:** Tunhe Zhou, Yulia Dragunova, Zegni Triki

## Abstract

Lately, there has been an emphasis on the importance of studying inter-individual variation in animal behaviour and cognition and understanding its underlying mechanisms. What was once considered mere noise around population mean can be explained by individual characteristics such as brain morphology and functionality. However, logistical limitations can be faced when studying the brain, especially for research involving wild animals, such as dealing with small sample sizes and time-consuming methods. Here, we combined an efficient and accurate method using X-ray micro-tomography and deep-learning (DL) segmentation to estimate the volume of six main brain areas of wild lizards, *Podarcis bocagei*: olfactory bulbs, telencephalon, diencephalon, midbrain, cerebellum and brain stem. Through quantitative comparison, we show that a sufficient deep-learning neural network can be trained with as few as five data sets. From this, we applied the trained deep-learning algorithm to obtain volume data of the six brain regions from 29 brains of *Podarcis bocagei*. We provide a detailed protocol for our methods, including sample preparation, X-ray tomography, and 3D volumetric segmentation. Our work is open-access and freely available, with the potential to benefit researchers in various fields, such as animal physiology, biomedical studies, and computer sciences.

## Introduction

There has been a growing interest in studying individual variations in behaviour and cognition and understanding their underlying neural correlates [1-4]. Understanding intraspecific variation in the brain and linking it to the ecology of the individual, like social group size, social dominance hierarchy, foraging behaviour, mating strategies, and predator avoidance [4-8] among other selective pressures, can help us to understand brain development and ultimately brain evolution. However, detecting individual variation in brain morphology can be difficult, especially in the endotherms [9]. In contrast, ectotherms with continuous adult neurogenesis allow for greater neural plasticity [10] and enable the capture of small-scale variation. Although fish brain morphology has been extensively researched in ectotherms due to their high plasticity, lizards possess comparable brain plasticity but are often overlooked.

Lizards can show complex cognitive abilities, such as flexibility and problem-solving, and possess a rich behavioural repertoire [11, 12]. Their geographic widespread distribution and occupancy of a wide range of ecological niches [12], in addition to their plastic brains, make them ideal for addressing questions about how their brain morphology shapes behaviour and cognition. For example, the wall lizards (*Podarcis*) are suitable study organisms for exploring individual variations in cognition and brain morphology that complement the variation in behaviour and physiology. Various behavioural studies have been conducted on *Podarcis*. For example, researchers have been interested in studying the fighting ability among different colour morphs of *Podarcis muralis* [13], as well as social learning in *Podarcis sicula*. However, few studies have linked behavioural differences to brain structures or functions [14].

There are, nevertheless, several limitations to conduct research aiming at linking brains to behaviour and cognition, especially in wild animals. Studying wild animals and obtaining decent sample sizes can prove challenging [15]. We often tend to maximise the amount of information and data that can be collected from wild specimens to compensate for small sample sizes but also for animal welfare purposes [16]. As part of increasing the amount and quality of data collected from wild specimens, using advanced technologies can prove extremely useful. For instance, using imagery methods, such as magnetic resonance imagery (MRI) and X-ray micro-tomography (microCT) to obtain 3D brain morphometric information for brain morphology enhances data quality and makes it more likely to detect small-scale inter-individual variation [4, 17]. These techniques generate high-resolution neuroanatomical images of the whole brain, surpassing the limitations of traditional histological methods that are both time-consuming and tissue-destructive [18-20]. To our knowledge, there is only one study on the lizard brain, the tawny dragon (*Ctenophorus decresii*), which has used the MRI technique [21]. While the study provides a reference brain atlas for the species, it employs manual segmentation that is challenging to reproduce. As part of advancing open science and improving the repeatability of the findings [3], we need accessible scripts and codes to recreate brain segmentations.

Here, we provide a detailed step-by-step protocol for extracting 3D brain morphometric data using X-ray microCT from a common lizard, *Podarcis bocagei*. To increase efficiency and reduce time-consuming sample preparation, we prepared whole heads instead of dissecting out the brains. This can also avoid potential damage during dissection, embedding and sectioning as in traditional histology. Using microCT 3D data and deep-learning (DL) neural network [22, 23], we segmented anatomically six distinct major brain regions: olfactory bulbs, telencephalon, diencephalon, midbrain, cerebellum and brain stem [24] of 29 specimens.

Deep learning (DL) segmentation for 3D images is fast growing in medical imaging [25], where the training data size is normally in the range of a hundred [26]. It is uncommon to have such big data quantity for wild animal studies. However, our study demonstrated that DL segmentation could provide satisfying results with as few as five training sets, which has the potential to significantly save manual segmentation work for smaller-scale studies in the biomedical field.

## Results and Discussion

### Virtual histology and 3D model from X-ray micro-tomography (microCT)

Compared to traditional histology, microCT is able to provide isotropic resolution in three dimensions in space, providing more detailed information in the sectioning dimension and allowing virtual histology in any direction from the same dataset. This significantly reduces manual work in sample sectioning and the number of animals. Examples of virtual histology sections of one brain from coronal, sagittal and horizontal directions are given in **Figure 1** and 2. Some parts of the brain regions are marked with references from [27-29] with the abbreviations reported in **Table 1**.

**Table 1.**
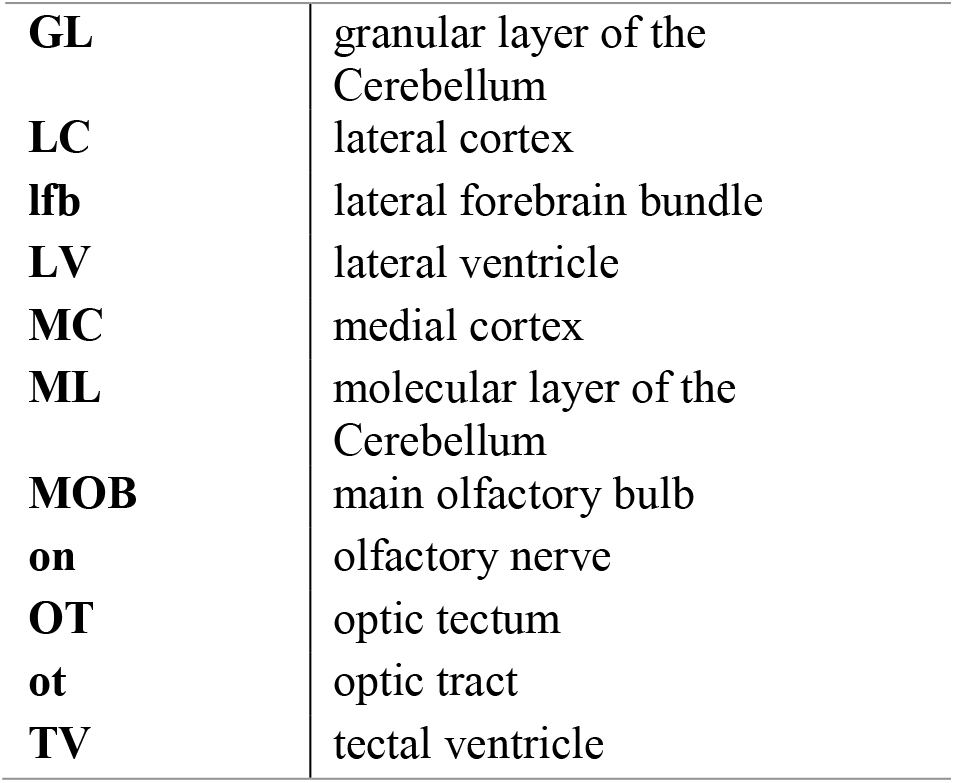
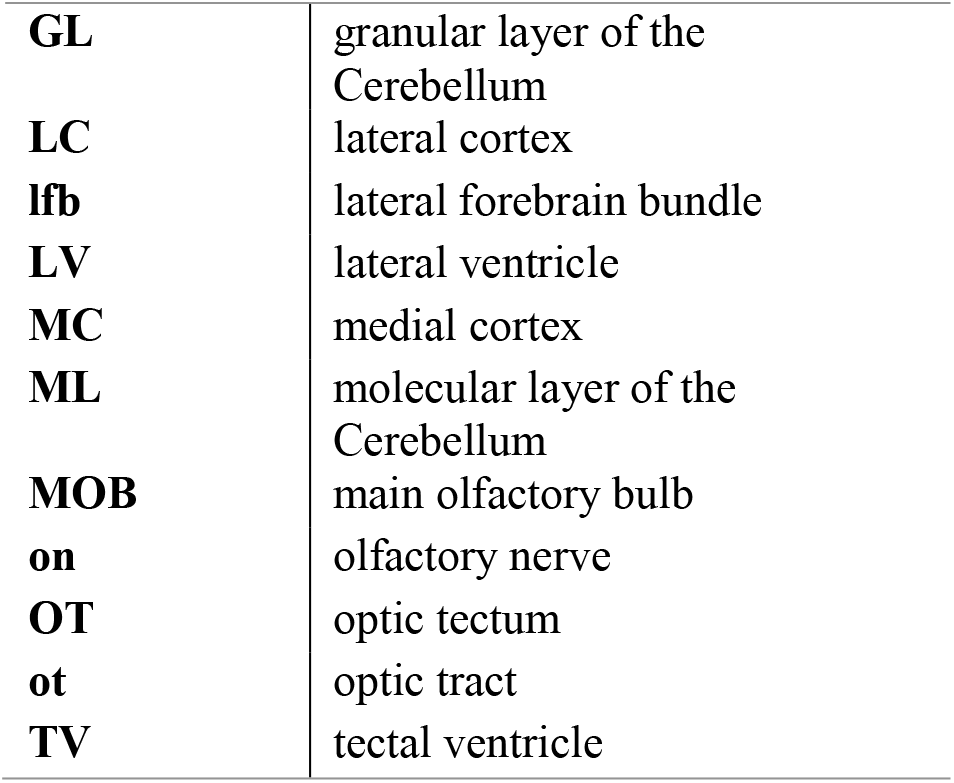
Abbreviations of brain regions.

**Figure 1.**
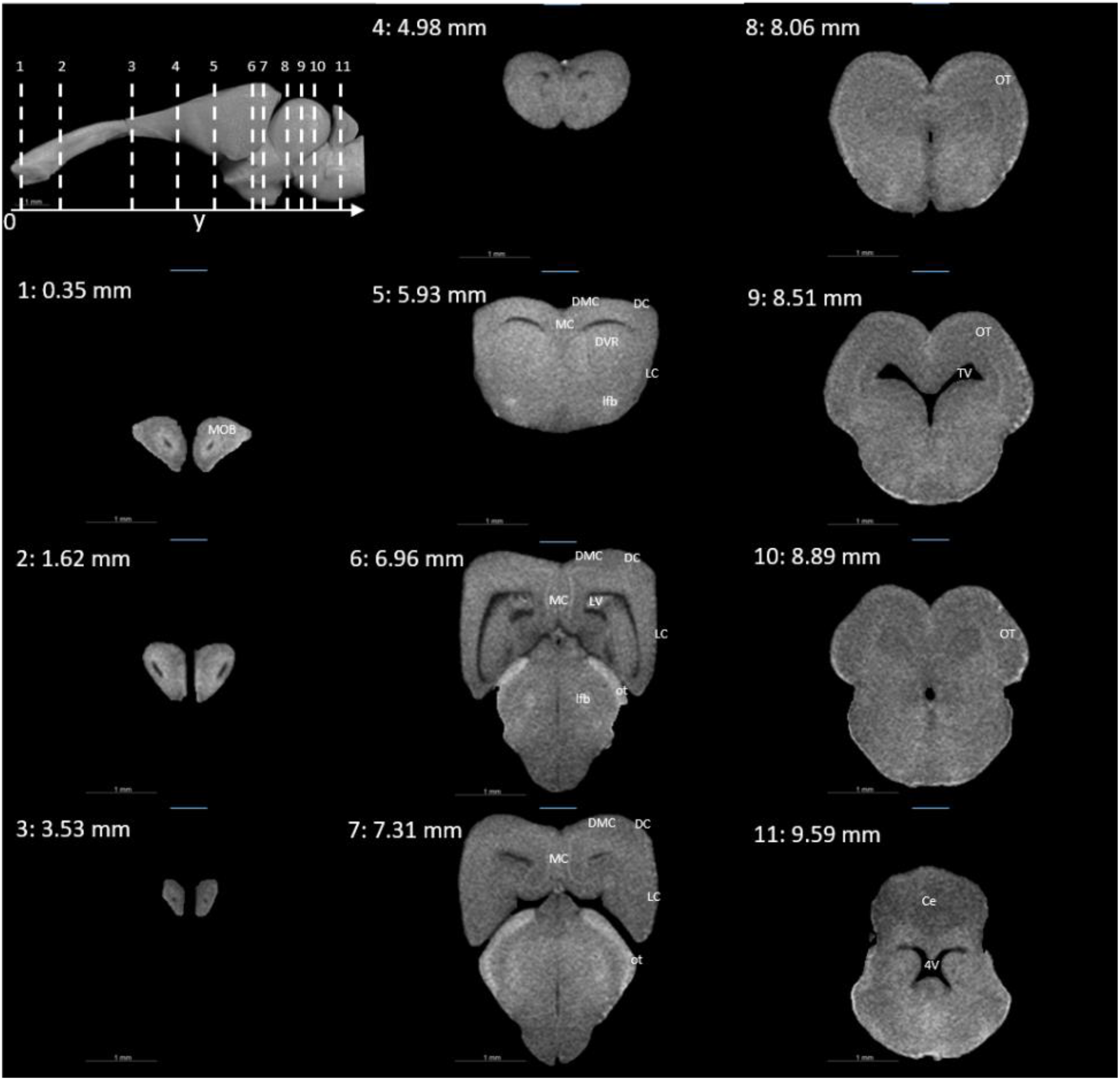
Coronal slices at positions indicated in the upper left subfigure. The abbreviations of the brain parts are listed in **Table 1**.

**Figure 2.**
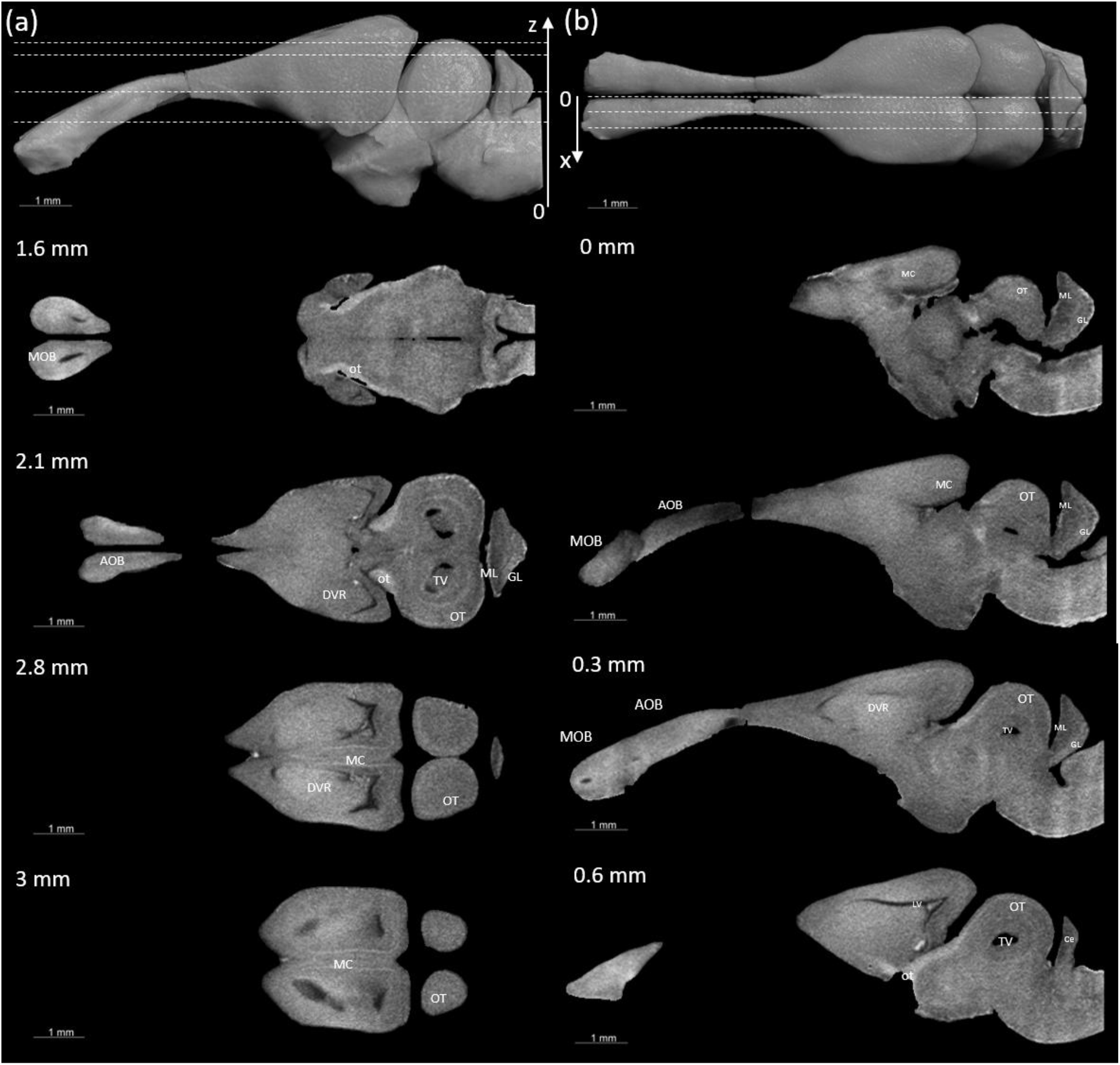
Examples of (a) horizontal and (b) sagittal slices from virtual histology at positions indicated in the upper subfigure. The abbreviations of the brain parts are listed in **Table 1**.

The brain virtual sections in **Figure 1** and 2 were generated with a mask from segmentation without dissection of the brain. The microCT images contain the whole head with detailed information about other tissues besides the brain. In comparison to magnetic resonance images (MRI), X-ray microCT provides higher contrast of bones and muscle tissue, and overall it yields higher resolution [27]. For example, in **Figure 3**(a) the olfactory nerves can be identified and we could measure the diameter to be about 26 μm as shown in the inset. In **Figure 3**(b), we showed another example of a horizontal slice demonstrating the possibility of using these images to study the visual system, providing complete 3D structures without the risk of deformation during dissection. It is important to have accurate 3D modelling to simulate the optics property and visual system of human [30] and animals [31], to reveal the resolution, focal length and field of view of the species to understand how they see the world. Our dataset can provide additional information for other studies on the species, for example, tooth development and skull structure as demonstrated in the cross sections in **Figure 3**(c,d) [32-34].

**Figure 3.**
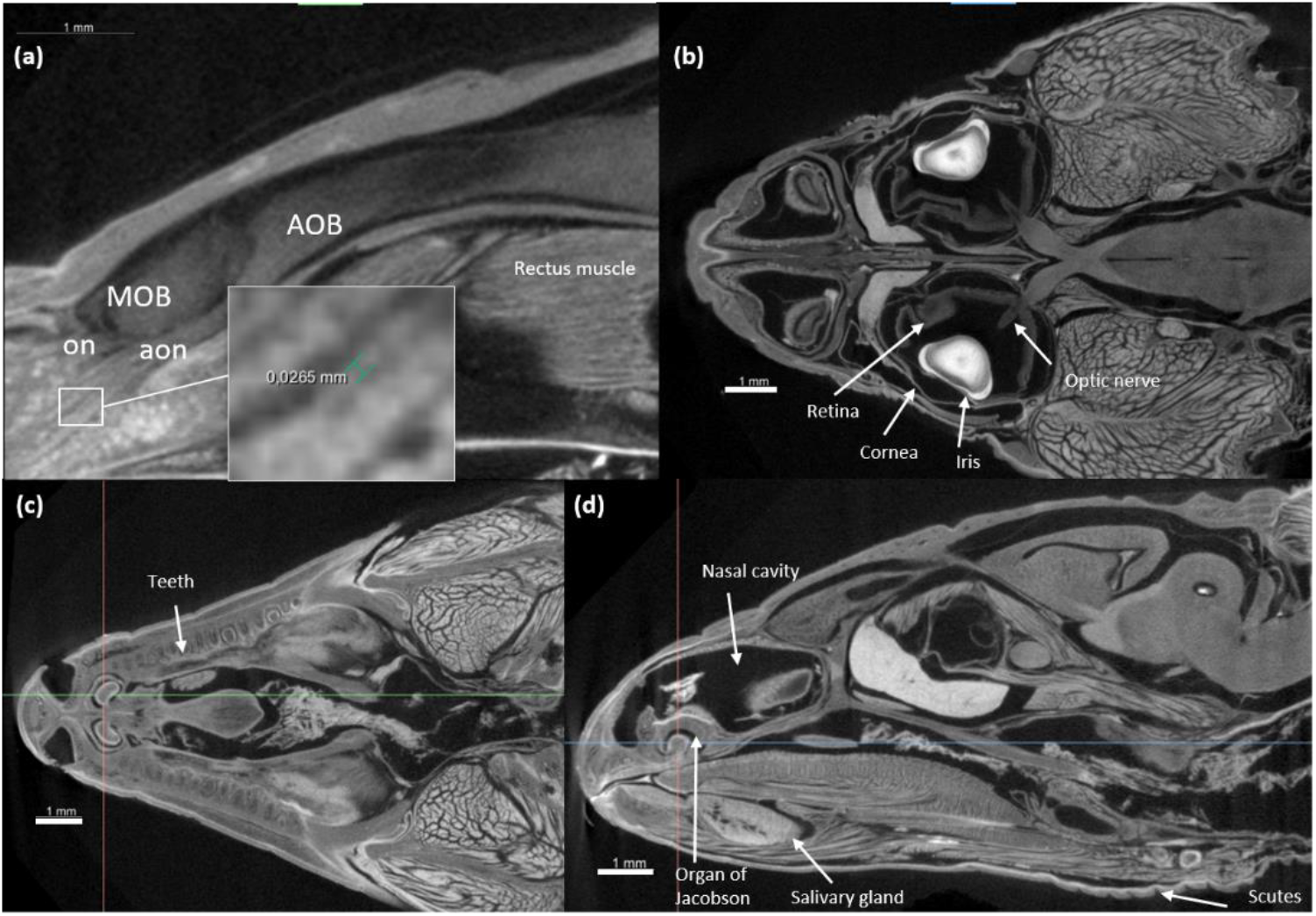
(a) Zoom in of a sagittal section on the olfactory system, showing the olfactory nerves (on) and accessory olfactory nerves (aon). (b) A horizontal section showing how the optic nerve connects the brain and the retina, as well as the cornea and iris of the eyes. (c, d) Transversal and sagittal cross sections of the head showing structures such as the teeth, salivary gland, the organ of Jacobson in the olfaction system.

### Brain region segmentation

We segmented six main brain regions from 29 specimens, namely, the olfactory bulb, telencephalon, diencephalon, midbrain, cerebellum and brainstem (see **Figure 4**). The detailed procedure of segmentation is reported in the Materials and Methods section. The volumes of the segmented brain regions of all samples were calculated from the number of voxels of each label multiplied by the voxel size. The average values are plotted in **Figure 5**, with the standard deviation as the error bar. Despite inter-individual variation in brain size, the variation in estimating region sizes was consistent and ranged between 14 and 19%.

**Figure 4.**
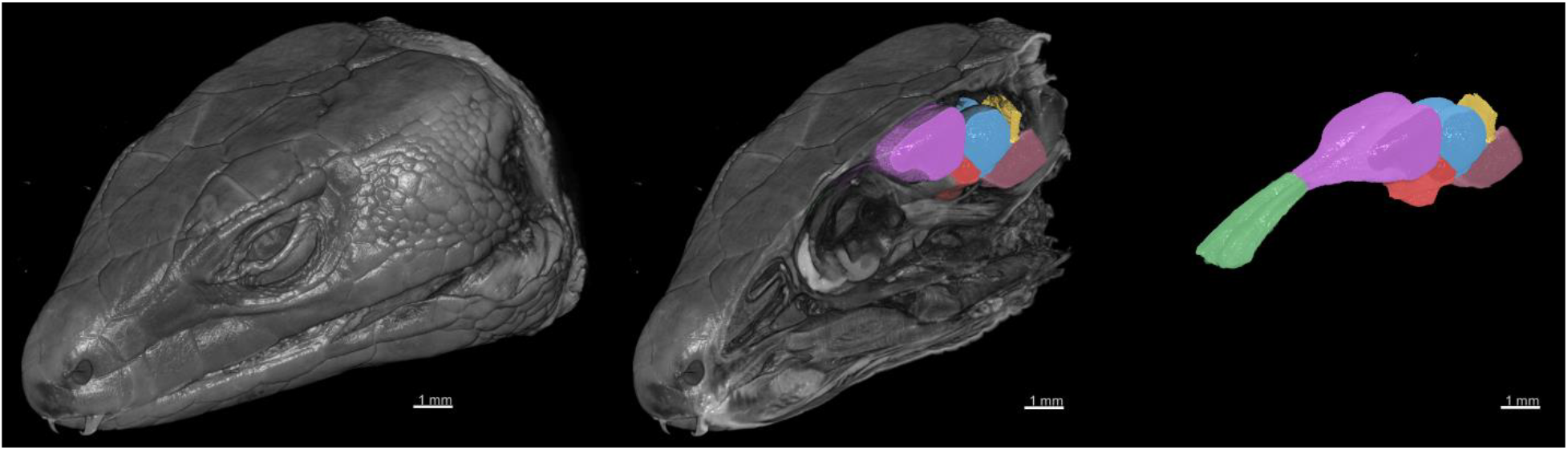
3D rendering of the X-ray microCT images showing the external and internal of the head, and the main parts of the brain. The brain regions are: the olfactory bulb (green), telencephalon (purple), diencephalon (red), midbrain (blue), cerebellum (yellow) and brainstem (pink).

**Figure 5.**
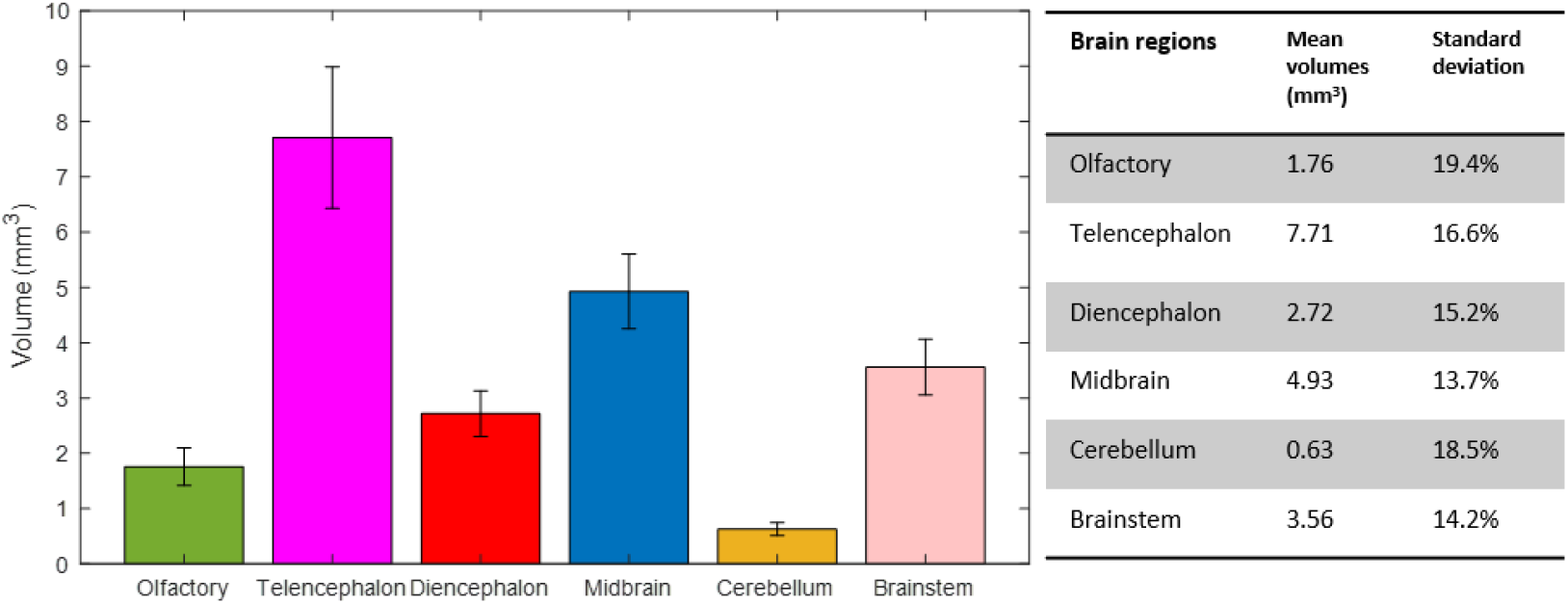
The volumes of the six brain regions of the 29 specimens were measured from the segmentation. The error bars show the standard deviation, and the bars show the mean value of the volumes.

Two deep learning algorithms were compared in this study for segmentation, namely Biomedisa and AIMOS [22, 35]. They were chosen as two of the most user-friendly open-source software, which is practically the most important factor for researchers in other fields to utilize the method in an efficient way with an easy installation and a short learning curve.

All three planes, coronal, sagittal and horizontal, were compared using Biomedisa in separate training and testing (see **Figure 6**a). For each of the planes, 1, 3, 5, 7, 9, and 11 images were used for training in 6 different runs and each trained neural network was used to predict 4 test image stacks. The predicted segmentations were evaluated using the average dice score (DSC) [36] and the average relative error of brain region volumes (RE). It can be seen in **Figure 6** (a) that as few as 3 samples could provide sufficient accuracy with DSC >0.9 using Biomedisa, and the three orientation planes resulted in no significant difference in DSC and RE when the training set is more than 3.

**Figure 6.**
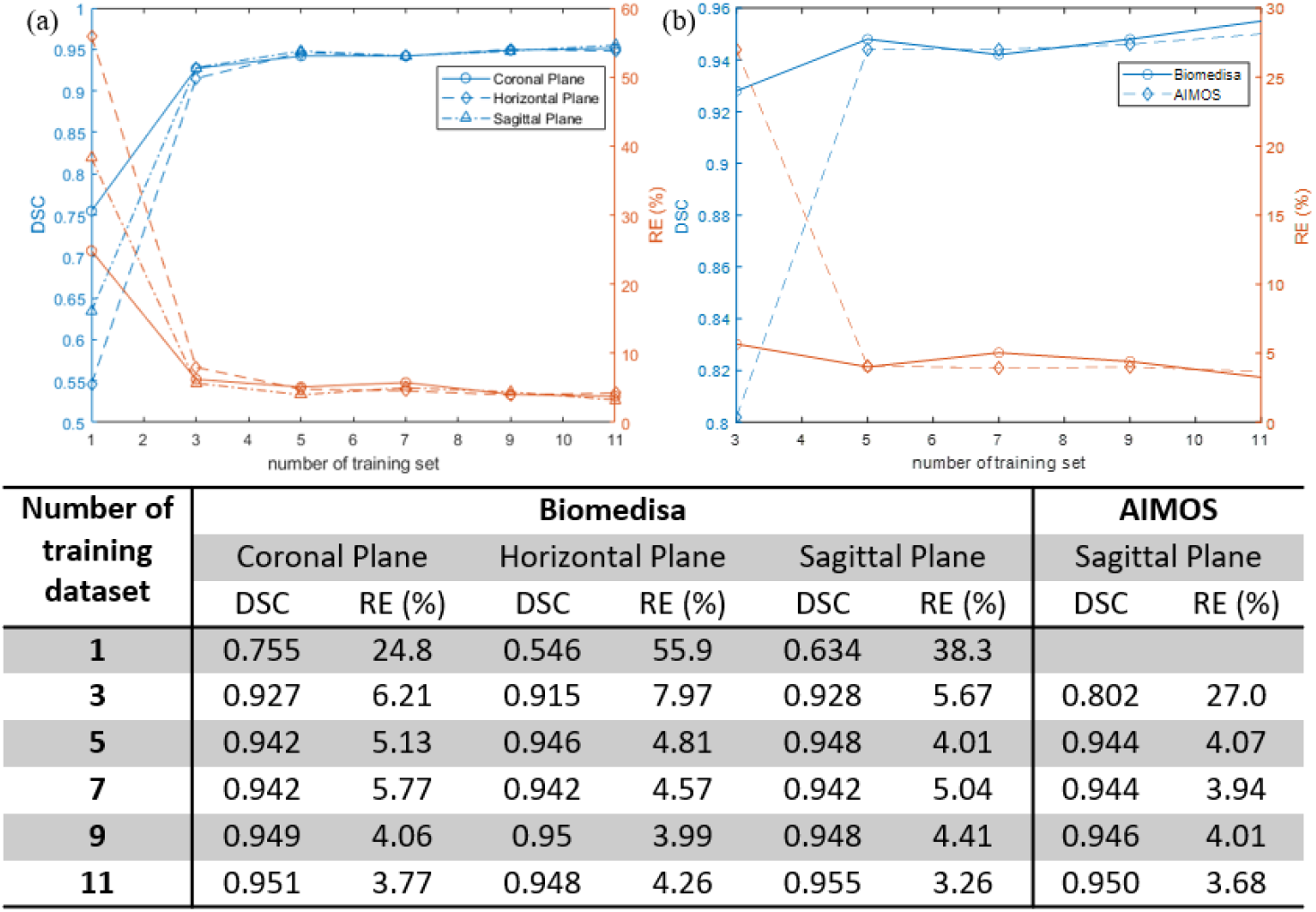
(a) Evaluation of predictions in the coronal plane using average dice score (DSC) and average relative error (RE) between the 6 brain regions as a function of number of training images. The resulting values are an average of 4 predicted segmentations. (b) Comparison of DSC and RE of Biomedisa and AIMOS in sagittal plane. The values of the plots are listed in the table [37].

In **Figure 6**(b) the performance of Biomedisa and AIMOS were also compared, and both were trained on the sagittal planes. As AIMOS requires 2 datasets from the labelled data for validation, the input data size cannot be smaller than 3. The comparison of the two algorithms reveals a significant difference for small data sizes due to similar factors. However, when the input data size is more than 5, this difference in performance disappears. We can conclude that for both algorithms, it is enough to use 5 labelled datasets to achieve an optimal dice value of over 0.94 and a quantitative difference in volume measurement of around 4% compared to manual segmentation.

In conclusion, 29 specimens of common lizard *Podarcis bocagei* were scanned using X-ray microCT to generate 3D images with internal structures in high resolution. Six brain regions, olfactory bulbs, telencephalon, diencephalon, midbrain, cerebellum and brain stem, were segmented, and their volumes were measured. Two DL segmentation algorithms were applied and compared quantitatively. The results have proven that as few as five training datasets (five samples) were sufficient for both algorithms. To verify the consistency of the DL segmentation with respect to the orientation of the sections, all three views, namely coronal, sagittal and horizontal slices, were compared in training and prediction accuracy, and no significant difference was detected. The methods described in this study offer an efficient protocol for achieving 3D imaging acquisition and volume measurement with minimal manual work and high accuracy. Accessing large brain specimens for ecology and evolution studies of vertebrates is often challenging. The approach we propose can help address some of these limitations, especially in studies on wild vertebrates with a small number of specimens, such as in our case with 29 lizards. Our methods achieve the objective of obtaining high-quality volumetric data within a short time frame and from a small sample size.

We hope that this protocol will be of helpful use to researchers from various fields beyond ecology and evolution, to obtain faster and more reliable (and repeatable) brain segmentation. Moreover, we make our 3D microCT data, as well as the labels and trained neural network, freely accessible. The shared 3D data of the whole head may benefit other research groups interested in head anatomical details beyond the brain, such as eyes, teeth, tongue, jaw bones and jaw muscles. Computer scientists could also benefit from accessing our data and code to develop more efficient algorithms for different tasks. Future work can focus more on improving the deep learning algorithms, combined with data augmentation, to push the limit of training data size to be even smaller and bring manual annotation efforts to a minimum.

## Materials and Methods

### Animals

This study used 29 male lizards (*Podarcis bocagei*) with a body mass of 4.38 ± 0.70 g (mean ± SD). They were captured in May 2021 around the Research Center in Biodiversity & Genetic Resources (CIBIO) campus, at the University of Porto, in Portugal. The animals were first used in another study to test their behaviour and cognitive abilities. Upon finishing the tests, all animals were euthanised with an anaesthetic intramuscular injection of Zoletil (10 mg/kg), followed by an intraperitoneal injection of sodium pentobarbital (80 mg/kg). The whole heads of the lizards were placed in a fixative solution composed of 4% paraformaldehyde for 10 days before being rinsed and stored in PBS. Then, the samples were shipped to Stockholm University for brain morphology analyses.

### Sample preparation

The lizard head samples were stained with Phosphotungstic acid (PTA) to enhance the tissue contrast for X-ray imaging. The staining protocol was adapted from Lesciotto et al. [38] to our samples for optimised scans. The samples have been already fixed in 4% PFA and rinsed and stored in PBS.

#### Dehydration process

30% ethanol in PBS 1 day;

50% ethanol in PBS 1 day;

70% ethanol in PBS 1 day;

one hour in a solution with a ratio of 4:4:3 volumes of ethanol, methanol, and H2O

80% methanol in ultrapure water for 1 hour;

90% methanol in ultrapure water for 1 hour.

#### PTA staining

Following dehydration, samples were immersed in 0.7% PTA in 90% methanol in ultrapure water.

During the staining, we were able to check the staining procedure several times within a few weeks until the whole brain was stained. For practicality, this step did not need to take the samples out of the PTA staining solution. We believe this would save the researcher significant time when checking the staining quality, as the samples did not have to be washed at this stage. Not all samples reached optimal staining at the same time point; rather, samples with relatively larger heads required prolonged staining times. Overall, the samples needed between 23 and 30 days of staining to reach optimality. The samples remained in the solution during scanning.

### X-ray microCT scan

The samples were scanned using Zeiss Xradia Versa 520 at Stockholm University Brain Imaging Centre, with the samples being stabilised inside 5mL plastic tubes and immersed in the storage solution, as shown in Figure 7. The X-ray source was set to have a voltage of 100 kV and a power of 9 W. The 0.4x objective and a CCD camera were used coupled with a scintillator. The effective voxel size was 17.4 μm with the optical and geometrical magnification compensated. The scan consisted of 801 projections over 212 degrees with 1 s exposure time for each projection. In total, one scan took 36 min including reference images and readout time of the CCD camera. Autoloader was utilized to change samples automatically to minimize the manual work. The tomography reconstruction was done automatically with Zeiss Scout-and-Scan software right after the scans and the output was 16-bit gray value tiff stacks.

**Figure 7.**
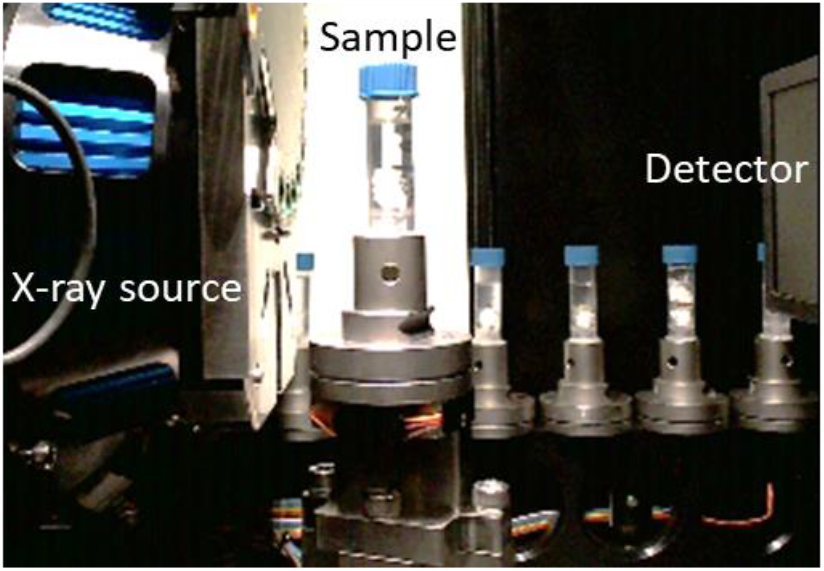
Experiment setup showing the X-ray scanner equipment and the samples prepared in a queue to be scanned automatically.

### Semi-manual segmentation of training datasets

It was not possible to manually place all samples in the same direction. For this reason, the first two steps in preparing the data were to align and crop all the images, as demonstrated in Figure 8(a). This was done using the software Dragonfly. This step may be overlooked but it was crucial for a consistent brain volume measurement, especially the brainstem part, and later for efficient neural network training.

**Figure 8.**
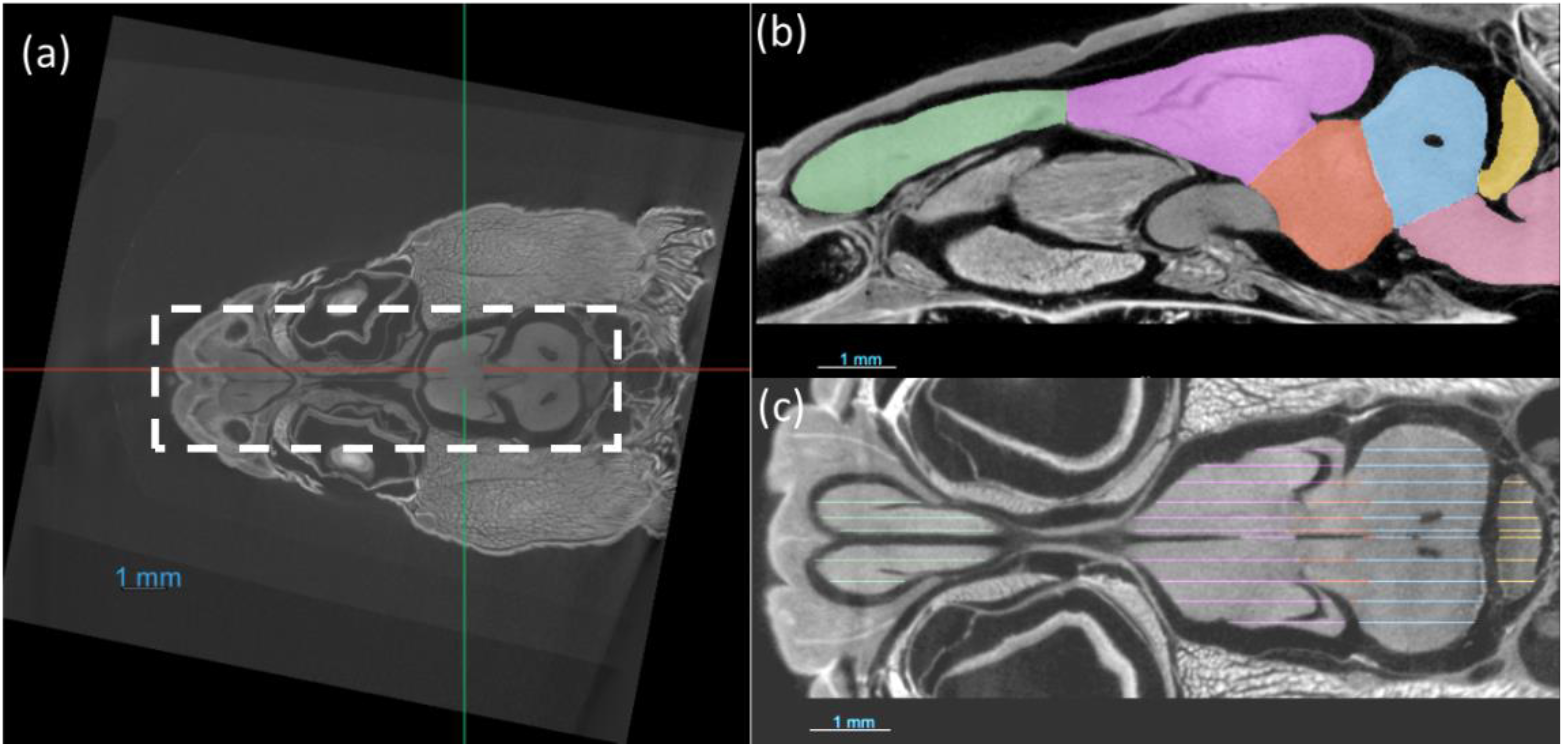
(a) The images are aligned and cropped to keep the brain for fast segmentation. (b,c) Demonstration of manual segmentation.

The training dataset was segmented semi-manually and consisted of three steps:

1) Manual segmentation of approximately 20-30 slices in Dragonfly (Figure 8 (b,c)).

2)Random walk interpolation in the Biomedisa [22, 39].

3) Check and correct manually.

### Deep-learning based segmentation

We tested and compared two DL algorithms, namely Biomedisa [22] and AIMOS [35]. Both of them are based on U-Net [40]. U-Net is a type of convolutional neural network (CNN) named for its unique U shape architecture, with an autoencoder with a skip connection. Compared to other CNN models, autoencoders delineate clear boundaries while maintaining a simpler model architecture. Additionally, the decoder in such networks is adept at forming distinct boundaries from the features it extracts. However, a significant challenge in using autoencoders is the potential over-simplification of images in the encoding phase [41]. To overcome this, linear skip connections are employed extensively. These connections enhance the accuracy of the segmentation maps by merging both elementary and complex features from different layers of the U-Net. Specifically, in U-Net, skip connections serve to directly transfer the detailed activation outputs from the encoder to their exact matching layers in the decoder, aiding in precise feature mapping.

### Biomedisa

We used the Biomedisa online app by preparing the images and labels in TAR format. Biomedisa follows 3D U-Net [42] and uses Keras with Tensorflow as framework [43]. Before training the network, tiff images were scaled and normalized, and 3D patches of a size of 64 × 64 × 64 voxels of the volumetric images were used as input. For training the network, we used the default parameter settings as shown in Table 2. The training time for using different numbers of datasets is listed in Table **3**. The rest of the dataset was predicted using the trained neural network where each took less than 1 min.

**Table 2.** Neural network training parameters Biomedisa.

|  |  |
| --- | --- |
| Network architecture | 32-64-128-256-512-1024 |
| Number of epochs | 200 |
| Batch size | 24 |
| Stride size | 32 |
| Image Scale | 256*256*256 |

**Table 3.** Training time of different dataset numbers using Biomedisa on sagittal planes [37].

| Number of training sets | Time (min) |
| --- | --- |
| 1 | 55 |
| 3 | 150 |
| 5 | 248 |
| 7 | 341 |
| 9 | 430 |
| 11 | 540 |

**Table 4.** The default neural network training parameters in AIMOS.

|  |  |
| --- | --- |
| Network architecture | 32-64-128-256-512-768 |
| Number of epochs | 30 |
| Batch size | 32 |
| Image Scale | 256*256 |

#### AIMOS

We also compared another user-friendly software AIMOS [35], using Python on a local computer and PyTorch as framework [44]. AIMOS uses an architecture similar to U-Net with 6 levels of encoding and decoding blocks. AIMOS has pre-trained networks available for microCT scans making the training time potentially shorter. In AIMOS, the data was split into a training for model weight optimization, a validation set for hyper-parameter optimization, and a test set for evaluation. The pipeline requires at least two extra datasets, compared to Biomedisa, where the validation ratio could be set to 0. This significant difference in demand minimizes manual labelling for small-scale studies like this project.

### Evaluation metrics

The dice score or dice similarity coefficient for each brain region was calculated using Eq. (1),

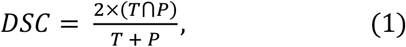

where T is the volume outlined by the manually segmented ground truth, and P is the volume outlined by the predicted segmentation [36]. The dice score is given between 0 and 1, where 1 indicates a perfect prediction. The dice score was calculated for each predicted brain region separately and then averaged to obtain the dice score for the whole sample.

One important value to measure in brain morphology analysis is the volume of the brain regions. It is, therefore, relevant to compare the accuracy of the volumes of the segmentation predictions generated from the deep learning algorithm. The average relative error was calculated from Eq. (2) as

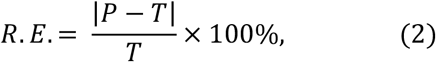

where P is the volume of the prediction, and T is the ground truth volume. The relative measurement error was calculated separately for all the brain regions in one sample and then averaged in order to obtain the relative error for each sample.

## Acknowledgements

We are grateful to Lekshmi Sreelatha, Zbyszek Boratynski, Miguel Carretero (and his lab), Bárbara Bastos, and Philipp Lehmann for generously providing us with the lizard samples and for their efforts in organising the logistics of animal capture, handling, sampling, and shipping. We also thank Anna Burvall for her advice on the segmentation part.

This work was financially supported by the Swiss National Science Foundation [grant numbers P400PB_199286 and PZ00P3_209020 to ZT] and a research grant from The Association for the Study of Animal Behaviour (ASAB) to ZT. Brain data acquisition was supported by a grant from the Stockholm University Brain Imaging Centre [grant number SU FV-5.1.2-1035-15].

## Ethics

Animals were provided under ethics permit no. 873-876/2021/CAPT granted to Miguel Carretero by Instituto de Conservação da Natureza e das Florestas (ICNF), Portugal. Animal capture, handling and euthanasia were conducted according to the regulations of the University of Porto.

## Data availability

The microCT data of the lizard brains and labels, as well as one example of the raw microCT data of a whole lizard head are shared on Figshare (DOI:10.17045/sthlmuni.26164570).

